# Abl kinase deficiency promotes AKT pathway activation and prostate cancer progression and metastasis

**DOI:** 10.1101/2020.05.19.104679

**Authors:** Melissa A Marchal, Devon L Moose, Afshin Varzavand, Destiney Taylor, James A Brown, Michael D Henry, Christopher S Stipp

## Abstract

Abl family kinases function as proto-oncogenes in various leukemias, and pro-tumor functions have been discovered for Abl kinases in solid tumors as well. However, a growing body of evidence indicates that Abl kinases can function to suppress tumor cell proliferation, motility, and *in vivo* tumor growth in some settings. To investigate the role of Abl kinases in prostate cancer, we generated Abl-deficient cells in a pre-clinical model of spontaneously metastatic, androgen-indifferent prostate cancer. Loss of Abl family kinase expression resulted in a highly aggressive, metastatic phenotype *in vivo* that was associated with AKT pathway activation, increased growth on 3D collagen matrix, and enhanced cell motility *in vitro*. Treatment of Abl kinase-expressing cells with the Abl kinase inhibitor imatinib phenocopied the malignant phenotypes observed in Abl-deficient tumor cells. In addition, inhibiting AKT pathway signaling abolished the increased 3D growth of Abl-deficient cells. Our data reveal that Abl family kinases can function as suppressors of prostate cancer progression and metastasis by restraining AKT signaling, a signaling pathway known to be associated with emergence of metastatic castration-resistant prostate cancer.

## Introduction

There will be an estimated 33,330 deaths due to prostate cancer in 2020 due largely to the emergence in patients of metastatic castration-resistant disease (mCRPC) [1]. Most mCRPC that emerges after first line androgen deprivation therapy remains androgen receptor (AR)-driven [2]. As a result, the potent next-generation AR pathway inhibitors (ARPIs) abiraterone and enzalutamide provide clinical benefits to a significant proportion of mCRPC patients [3, 4]. Nevertheless, the eventual emergence of mCRPC that resists all currently available ARPIs remains a major clinical challenge.

Mechanisms of resistance to ARPIs include AR amplification [5], AR splice variants and point mutations [6-9], and upregulation of the glucocorticoid receptor, which can promote transcription of AR target genes [10]. However, a subset of ARPI-resistant prostate cancers display basal, mesenchymal, squamous, neuroendrocrine (NE)-like, or AR/NE-double negative phenotypes [11-14]. These “AR-indifferent” tumors activate alternative pathways, including neuronal, cell adhesion, epithelial-mesenchymal transition (EMT), and stem cell pathways [12, 13]. One recent study found evidence of NE-like disease in 17% of 160 patients whose cancer had progressed on abiraterone or enzalutamide, a feature associated with shorter overall survival [15]. No targeted therapies exist for AR-indifferent mCRPC, thus there is an urgent need to identify clinically actionable targets that promote proliferation and survival of mCRPC cells that no longer rely on AR signaling.

Hyperactivation of PI3K/AKT signaling due to PTEN deficiency can promote the development of mCRPC [16, 17], which may be linked mechanistically to reciprocal inhibition between the AR and PI3K/AKT signaling pathways [18-20]. The emerging view is that the PI3K/AKT and AR signaling pathways mutually inhibit each other, and therapeutic inhibition of either pathway alone leads to compensatory activation of the other pathway. This insight led to a recent clinical trial using the AKT inhibitor ipatasertib in combination with abiraterone in patients with mCRPC, resulting in improved radiographic progression-free survival compared to abiraterone alone, particularly for PTEN-deficient tumors [21]. However, the presence or absence of AR-indifferent or NEPC-like disease was not evaluated in this trial, and the potential utility of targeting AKT signaling specifically in AR-indifferent mCRPC remains to be explored.

Although Abl kinases can have tumor promoting functions [22], a growing body of evidence indicates that Abl kinases can function to restrain malignant behavior in multiple cancers [23-32]. We previously identified Abl kinases as potential metastasis suppressors operating downstream of *α*3 integrin in a PTEN-deficient model of mesenchymal, AR-indifferent mCRPC [33]. However, the role of Abl kinases in mCRPC progression and dissemination *in vivo* was unknown, and the molecular mechanism by which Abl kinases can restrain malignant behaviors of mCRPC cells remained to be defined. We now find that Abl kinase-deficiency dramatically potentiates CRPC progression and metastasis *in vivo*, which corresponds to upregulation of AKT signaling and tumor cell growth on 3D matrix, and increased tumor cell motility. Our results reveal a novel mechanism by which Abl kinases function to inhibit tumor progression, and support the rationale of targeting AKT signaling in PTEN-deficient, AR-indifferent mCRPC.

## Materials and methods

### Cell lines and culture

GS689.Li cells are a metastatic variant of PC-3 prostate cancer cells, that were created by *in vivo* passaging of PC-3 prostate cancer cells in SCID mice [34], and subsequently reverified by STR analysis (IDEXX Bioresearch) [33]. TEM4-18 cells are a subpopulation of PC-3 prostate cancer cells selected for enhanced transendothelial migration *in vitro* [34]. DU-145 cells, LNCaP cells, VCaP cells, and 22Rv1 cells were obtained directly from ATCC. Growth base media and supplements were from Gibco ThermoFisher. GS689.Li cells, TEM4-18 cells, and DU-145 cells were cultured in DMEM:F12, LNCaP cells and 22Rv1 cells were cultured in RPMI, and VCaP cells were cultured in DMEM. Growth media were supplemented with glutamine, penicillin/streptomycin, non-essential amino acids, and 10% FBS (Atlanta Biologicals). All cell lines harbored a pQCXIN retroviral expression vector encoding luciferase.

### Antibodies and reagents

This study utilized antibodies from Cell Signaling Technology against FOXA2 (#8186), Rb (#9309), c-ABL (#2862), FAK (#3285), SRC (#2110), phospho-SRC Tyr416 (#6943), ERK (#4696), phospho-ERK (#9101), phospho-AKT Ser473 (#4060), phospho-AMPKα Thr172 (#2535), Cyclin D3 (#2936), phospho-S6 Ribosomal Protein Ser240/244 (#5364), and AXL (#8661). Also used were antibodies from BD Transduction against E-cadherin (#610181), phospho-FAK Tyr397 (#611806), and AKT (#610860), as well as *α*-tubulin (12G10, Developmental Studies Hybridoma Bank), phospho-AXL Tyr779 (MAB6965, R&D Systems), chromogranin A (NB100-79914, Novus Biologicals), and ARG (Arg 11, generous gift from Anthony Koleske, PhD, Yale University). Secondary antibodies used were goat-anti-mouse and goat-anti-rabbit antibodies conjugated to Alexa 790, 680, 594, or 488 (Invitrogen). Other reagents were D-luciferin (Gold Biotechnology), rat tail collagen I (Corning & Advanced Biomatrix), imatinib (Cayman Chemical), and MK-2206 and R428 (Selleckchem).

### RNA interference

The specificity and efficacy of ABL and ARG knockdown was previously validated by multiple independent RNAi targeting vectors [33]. We used the most effective RNAi vectors from our previous study to create Abl (ABL KD) and Arg (ARG KD) single and double knockdown (ABL/ARG KD) GS689.Li cells. The Abl sh3 retroviral construct has a pZIP-mCMV-ZsGreen backbone (Transomics Technologies) and the Arg sh2 retroviral construct has a pSIREN RetroQ vector backbone (Clontech).

All four cell lines were carefully matched by passage number and contained both vector backbones. Thus, vector controls (NT/NT) contained both non-targeting shRNAs in the corresponding vector backbones, single knockdowns contained one targeting shRNA in one vector backbone and one non-targeting shRNA in the other vector backbone, and double knockdowns contained both targeting shRNAs in the corresponding vector backbones. Because each cell line in the set contained a pZIP-mCMV-ZsGreen vector, all cells were fluorescently labeled. Cells were maintained as stably transduced, polyclonal populations. The RNAi targeting sequences used for the present study were as follows: Abl sh3, 5’-GCAGTCATGAAAGAGATCAAA-3’ (ULTRA-3267084, Transomics); and Arg sh2, 5’-CCTCAAACTCGCAACAAAT-3’.

### Orthotopic prostate cancer model

All animal protocols were approved by the University of Iowa Institutional Animal Care and Use Committee (Approval #8031328). On the day of inoculation, 5 × 10^4^ NT/NT, ABL KD, ARG KD, and ABL/ARG KD GS689.Li cells were implanted in the left or right anterior lobe of the prostate of 10 SCID/NCr (BALB/C) mice/cell line. Bioluminescent imaging (BLI) was performed using an Ami X imaging system (Spectral Instruments Imaging) as described previously [33]. Upon sacrifice, livers, kidneys, and lungs were dissected for imaging of disseminated cells by fluorescence microscopy using an Olympus SZX12 stereomicroscope, as described [33]. GS689.Li cells from primary tumors were recovered by mincing with a sterile razor blade and digesting with 200 U/ml collagenase II (Worthington Biochemical Corporation) for 15 min at 37°C. Explanted cells were grown out under G418 to select for luciferase-positive tumor cells.

### Time-lapse motility assays

A total of 2.3 × 10^5^ cells were plated in serum-free medium [SFM; DMEM:F12, supplemented with 5 mg/mL BSA, glutamine, penicillin/streptomycin, non-essential amino acids, 25 mM HEPES pH 7.2, and insulin–transferrin–selenium (ITS)] on 35-mm dishes coated with 10 ug/mL rat tail collagen I. Time-lapse images were acquired as described previously [33], and migration speed and displacement were calculated using the mTrackJ plugin [35]. ImageJ software was used to measure the morphological characteristics of migrating cells (at t = 20 minutes) by making cell traces using the Freehand selections tool, followed by the Measure (Shape descriptors) command.

### Immunostaining

Cells were fixed with 10% formalin in HBSM (HEPES-buffered saline with 1 mM MgCl_2_), rinsed with Tris-buffered saline, then blocked and permeabilized with 0.1% NP-40 detergent in 10% goat serum prior to exposure to primary antibodies. Primary antibodies were detected with Alexa 488 or 594-labeled secondary antibodies and coverslips were rinsed and mounted in ProLong Gold with DAPI (ThermoFisher Scientific) for fluorescence microscopy.

### 3D collagen growth assays

Twenty-four-well plates were coated with 350 µL of rat tail collagen I (0.8 mg/mL in DMEM) per well and allowed to polymerize for 45 minutes at 37°C. A total of 1.5×10^4^ cells per well were seeded in 500 µL of SFM. After 7–10 days, cell number was quantified by BLI. Six wells per cell type per condition were quantified in each trial.

### Standard growth assays

A total of 1.5×10^4^ cells per well were plated in uncoated 24-well plates in 500 µL of standard growth medium. After 4–7 days, cell number was quantified by BLI. Six wells per cell type were quantified in each trial.

### Immunoblotting

Cells were rinsed twice with HBSM and lysed in Laemmli buffer. For analysis of cells growing in 3D conditions, cells growing on 0.8 mg/ml collagen I were rinsed twice with HBSM and then incubated two minutes at room temperature with Laemmli buffer. Lysate was collected from the surface of the 3D matrix by pipetting. Protein concentrations of lysates were normalized using the RED 660 Protein Assay (G Biosciences) prior to SDS-PAGE and immunoblotting. Blots were analyzed using a LiCOR Odyssey blot imager.

### Reverse phase protein array (RPPA)

Cells growing on 3D collagen for 4 days were lysed in 10% glycerol, 2% SDS, 0.0625 M Tris-HCL, pH 6.8, and 2.5% beta-mercaptoethanol, and lysate concentrations were adjusted to 1.0 ug/uL. Duplicate lysates of NT/NT and ABL/ARG KD cells were analyzed at the Functional Proteomics RPPA core facility (MD Anderson Cancer Center). Normalized linear protein levels for each of 304 signaling proteins included in the analysis were used to calculate the normalized linear fold change [(normalized linear value ABL/ARG KD)/(normalized linear value NT/NT)], and a cutoff was set for an average fold change of 1.2 or greater or 0.8 or less. A heatmap was generated using median centered log_2_ values from targets meeting the cutoff criteria, using the Heatmapper tool (http://www.heatmapper.ca/).

### Statistics

Statistical analyses were performed using GraphPad Prism Version: 7.02 (GraphPad Software, Inc.) Student’s t-test and ANOVA were employed for pairwise and multiple comparisons, respectively. When appropriate, Mann-Whitney Rank Sum and ANOVA on Rank (Kruskal-Wallis) tests were also used.

## Results

### GS689.Li cells are a model of mCRPC with mesenchymal and neuroendocrine-like features

We previously described the isolation of GS689.Li cells, a metastatic variant of the PC-3 prostate carcinoma [34]. As described by other groups [36], we found that cells recovered from the rare metastases that emerge from PC-3 xenografts were more metastatic upon re-injection [34]. Strikingly, these metastatic variants displayed loss of E-cadherin and upregulation of a mesenchymal gene expression program [34]. Since PC-3 cells have been reported to possess NE-like features [37, 38], and EMT and NE phenotypes are intertwined [13], we investigated that interconnection in GS689.Li cells. Both parental PC-3 and GS689.Li cells appeared positive for the NE marker chromogranin A (Fig. S1A, B), as previously reported for parental PC-3 cells [39]. The PC-3 parental cell population is p53-null [40], and dual loss of p53 and Rb can drive an NE-like phenotype [41, 42]. Nevertheless, both PC-3 and GS689.Li cells retain Rb expression (Fig. S1C, D), as previously described for the parental PC-3 cells [43]; however, PC-3 cells are PTEN-deficient [44], and loss of PTEN or activation of AKT can also contribute to an aggressive phenotype with neuroendocrine features [41, 45]. Recently, transcription factor FOXA2 has been described as a marker for a significant fraction of neuroendocrine-like prostate cancers [37]. We detected a FOXA2-positive subpopulation in the parental PC-3 cell line that corresponded largely to the E-cadherin-negative sub-population in double labeling experiments (Fig. S1E). Strikingly, 100% of the GS689.Li cells were E-cadherin-negative/FOXA2-positive, with strong nuclear FOXA2 localization (Fig. S1F). Collectively, these data reveal that parental PC-3 cells already express some features of NE-like CRPC, but that *in vivo* passaging selected for a metastatic, FOXA2-positive, mesenchymal subpopulation.

### Abl kinases function as suppressors of mCRPC tumor progression and metastasis

To investigate the role of the Abl family kinases in mCRPC tumor growth and metastasis, we depleted ABL1 (ABL KD), ABL2 (ARG KD) or both kinases (ABL/ARG KD) in GS689.Li cells. Luciferase/ZsGreen dual-labeled, Abl kinase-deficient and non-targeting control cells (NT/NT), were orthotopically implanted, and *in vivo* tumor progression was monitored weekly via BLI. By day 35 after implantation, mice harboring ABL KD cells and ABL/ARG KD cells displayed a markedly higher apparent tumor burden than mice harboring ARG KD or NT/NT control cells (Fig. 1A). Across the course of the experiment, Abl kinase-deficient tumors progressed more rapidly than control tumors (Fig. 1B). Loss of ABL had a large effect, but loss of both ABL and ARG had the largest effect, resulting in tumor burdens over an order of magnitude greater than NT/NT control tumors by day 35.

**Figure 1:**
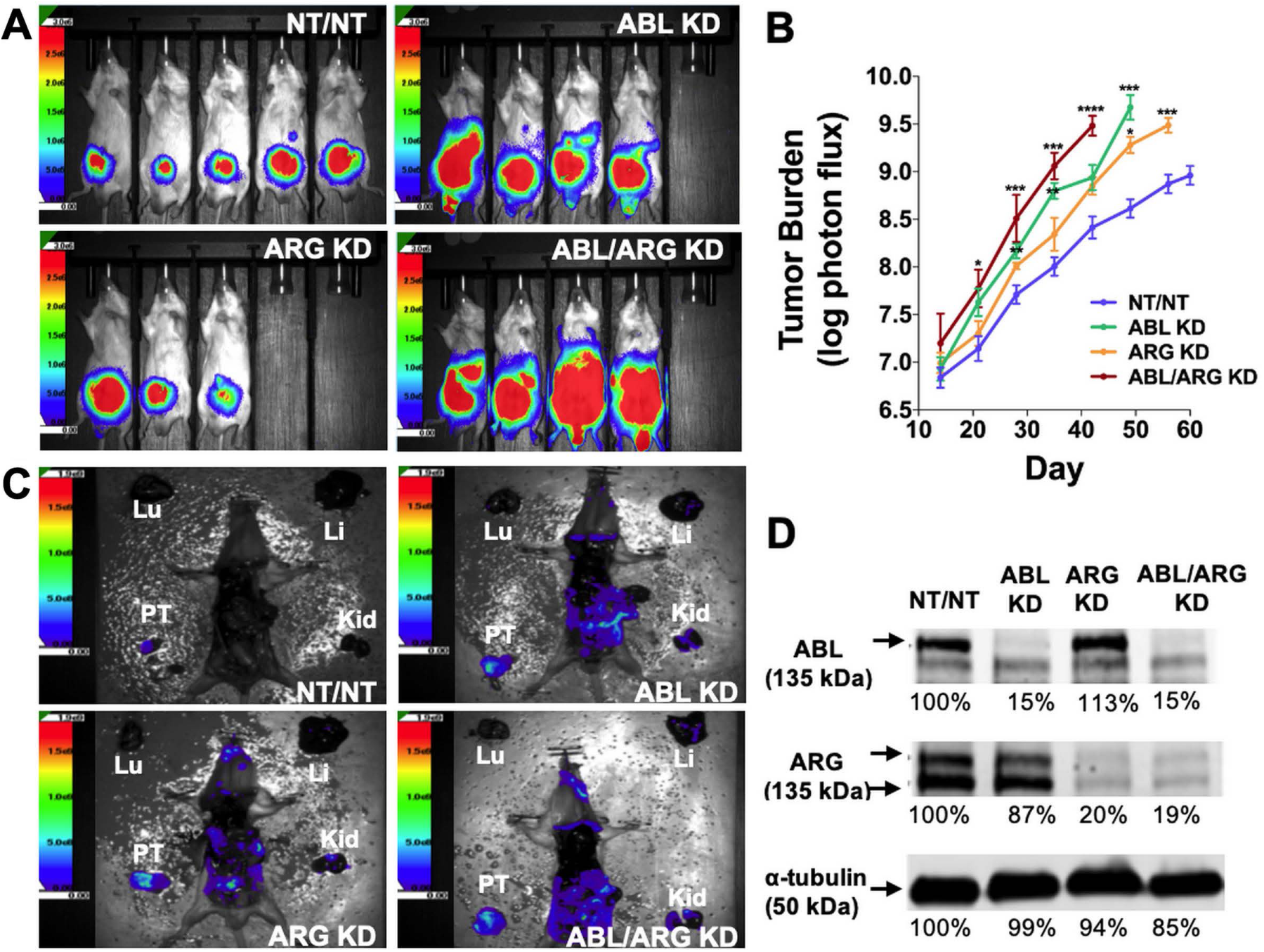
Loss of Abl family kinase expression promotes *in vivo* mCRPC tumor progression and metastatic dissemination. **(A)** Representative bioluminescence (BLI) images of orthotopic tumors in mice implanted with Abl family kinase-deficient (ABL KD, ARG KD, and ABL/ARG KD) and non-targeting (NT/NT) mCRPC cells at 35 days post-implantation. **(B)** Semi-log plot depicting average tumor burden (log of photon flux [photons/s]) over time (day) for mice bearing NT/NT (blue line), ABL KD (green line), ARG KD (orange line) and ABL/ARG KD (red line) tumors. Data points indicate mean +/- SEM for ≥ 7 mice/cell type. *, **, ***, and **** signify statistically significant p-values of < 0.05, < 0.01, < 0.001, and < 0.0001, respectively. Kruskal Wallis with Dunn’s multiple comparison test and Mann Whitney test at α = 0.05. **(C)** Representative images of BLI-assisted gross necropsies performed immediately after sacrifice for mice harboring NT/NT (sacrificed day 63), ABL KD (sacrificed day 52), ARG KD (sacrificed day 59), and ABL/ARG KD (sacrificed day 45) tumors. PT: primary tumor; Kid: kidney; Li: liver; Lu: lung. **(D)** Immunoblot analysis of ABL and ARG proteins in all mCRPC cell types prior to implantation. % values represent blot intensities normalized by an α-tubulin loading control and expressed as percent relative to NT/NT. Color scales in A. and C. indicate photons/sec/cm^2^/sr.

To attempt to control for effects of primary tumor size, we allowed mice in the different groups to age out for different times until their total tumor burdens approached a similar value, although the vector control group still had lower average tumor burden at the time of sacrifice. Shown in Fig. 1C are gross necropsy images of mice with NT/NT control, ABL KD, ARG KD, and ABL/ARG KD tumors. The apparent tumor burden of all four mice was ∼1 × 10^9^ photons/sec prior to sacrifice, yet the dissemination of Abl kinase-deficient tumor cells appeared qualitatively substantially increased compared to vector control tumor cells (Fig. 1C). Analysis of tumor cells explanted at the time of sacrifice revealed an 80-85% reduction in Abl and Arg proteins (Fig. S2), which was similar to what we observed in ABL and ARG KD cells prior to implantation (Fig. 1D), demonstrating that ABL and ARG knockdown was maintained in tumor cells *in vivo*.

We used GFP fluorescence microscopy to further investigate tumor cell dissemination to specific organs. ABL KD and ABL/ARG KD tumor cells both displayed significantly increased dissemination to kidney (primarily in the adjacent adrenal gland), liver, and lung compared to vector control cells (Fig. 2A-L, quantified in M-O). For ARG KD single knockdown cells, there was a trend towards increased kidney colonization, although it did not rise to the level of statistical significance in this analysis (Fig. 2M).

**Figure 2:**
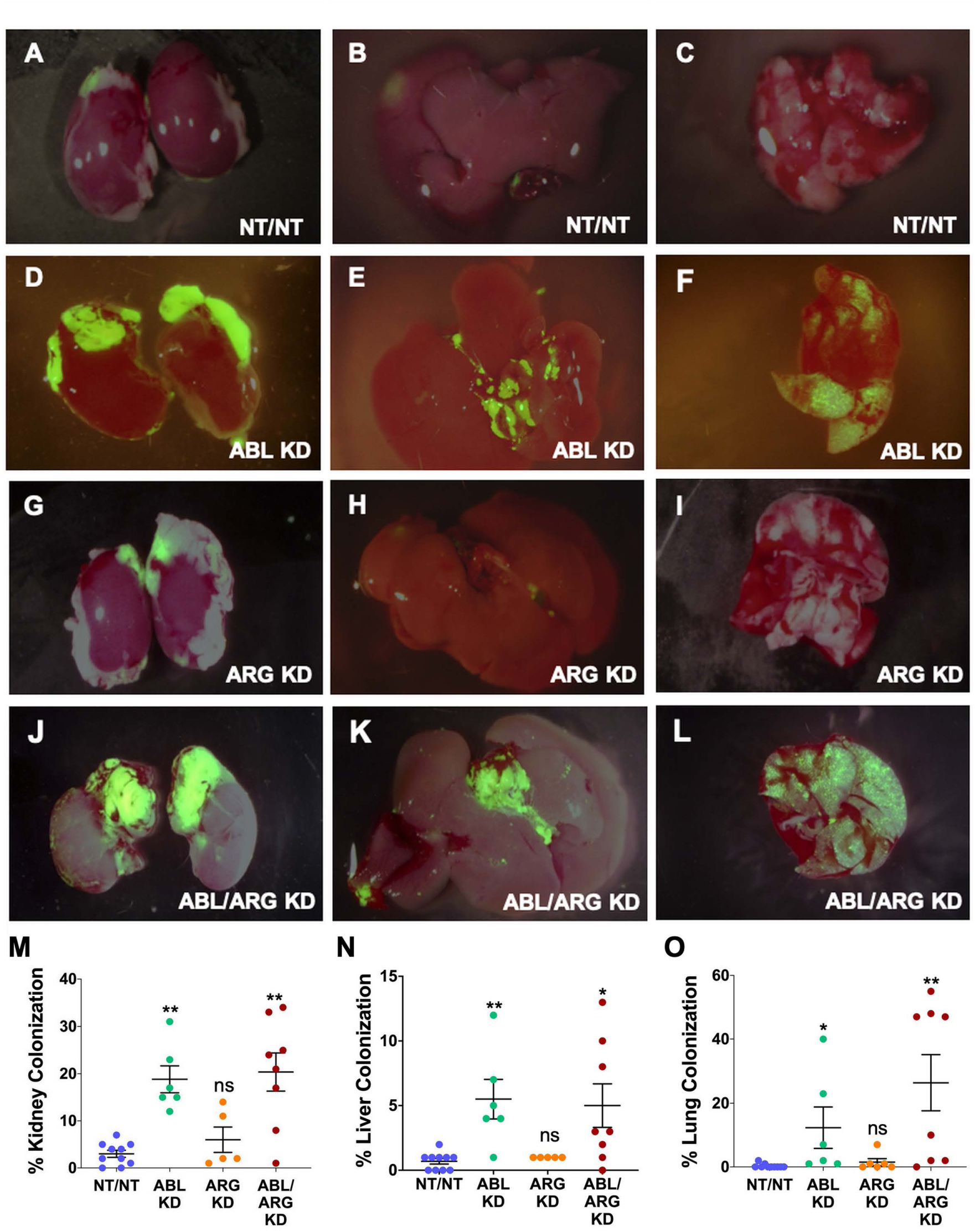
Abl kinase-deficient tumor cells exhibit enhanced metastatic colonization. Representative fluorescent micrographs of kidneys (1^st^ column), livers (2^nd^ column), and lungs (3^rd^ column) colonized by GFP-labeled NT/NT **(A, B, & C)**, ABL KD **(D, E, & F)**, ARG KD **(G, H, & I)**, and ABL/ARG KD **(J, K, & L)** mCRPC cells. Quantification of kidney **(M)**, liver **(N)**, and lung **(O)** colonization as the percentage of total surface area occupied by GFP-labeled mCRPC cells. Bars in M, N, & O indicate mean +/- SEM for ≥ 5 organs/cell type. * and ** in M, N, & O denote statistically significant p-values of < 0.05 and < 0.01, respectively, and n.s. stands for not significant. Kruskal-Wallis with Dunn’s multiple comparison test, α = 0.05.

Because it is difficult to fully account for the contribution of more rapid progression of the primary tumor to increased metastatic dissemination, we also compared the motility of Abl-deficient tumor cells to vector control cells via time lapse video microscopy. ABL/ARG KD cells displayed dramatically increased motility compared to vector control cells, as visualized by cell motility tracks (Fig. 3A), with ABL KD and ARG KD single knockdown cells displaying an intermediate phenotype. Quantification revealed a striking ∼2-fold increase in migration speed and net displacement of ABL/ARG KD cells compared to vector control cells, with ARG single knockdown producing a larger effect than ABL single knockdown on cell motility. (Fig. 3B, C). The enhanced motility of Abl kinase-deficient cells was associated with a morphological change from round, spread vector control cells to highly polarized Abl kinase-deficient cells, with prominent leading edge lamellipodia (Fig. S3A-D). This morphological change is quantitatively captured as a significant increase in the aspect ratio of ARG KD and ABL/ARG KD cells (Fig. S3E), and a significant reduction in roundness (Fig. S3F).

**Figure 3:**
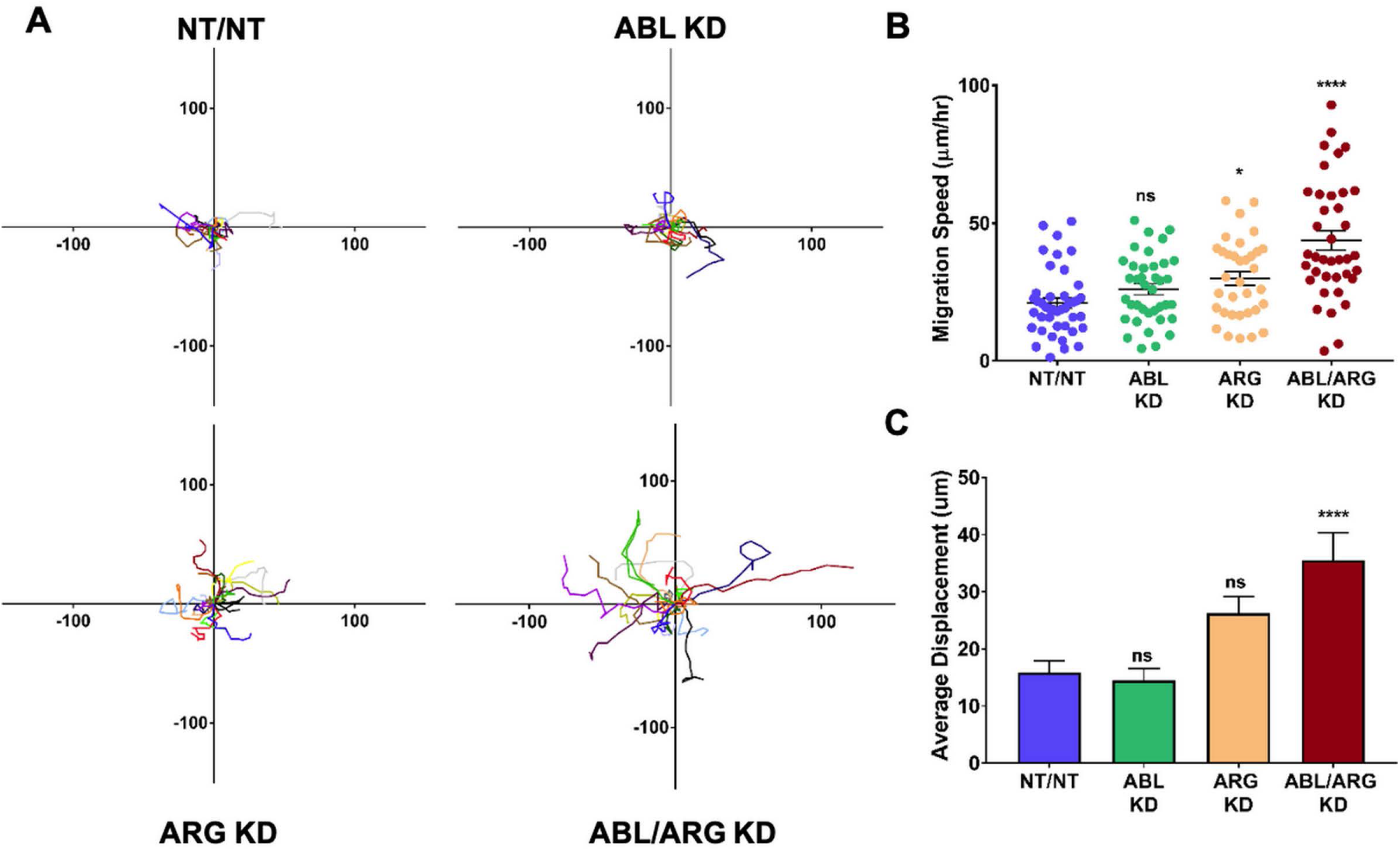
The Abl family kinases act to constrain the motility phenotypes of mCRPC cells. Abl family kinase-deficient (ABL KD, ARG KD, and ABL/ARG KD) and non-targeting (NT/NT) mCRPC cells were plated on 2D collagen I and had their migration monitored via time-lapse microcopy for 90 minutes. **(A)** Motility tracks; n = 20 tracks/cell type. X/Y axes in um, 0 represents point of origin. **(B)** Graph of average migration speed (um/hr.), n = > 34 cells/cell type. **(C)** Graph of average displacement (um) from point of origin, n = > 34 cells/cell type. Bars in B & C indicate mean +/- SEM. * and **** in B & C indicate statistically significant p-values of < 0.05 and < 0.0001, respectively, and n.s. stands for not significant. One-way ANOVA w/ Dunnett’s multiple comparison, α = 0.05.

Taken together, these data revealed Abl kinases can function as suppressors of progression and metastatic dissemination of mCRPC cells, restraining tumor growth, metastatic colonization, and cell motility. Loss of ABL had a larger effect on tumor growth, while loss of ARG had a larger effect on cell motility. These observations may help to explain why loss of both kinases produced the most aggressive phenotype in the *in vivo* tumor progression model.

### Loss of Abl kinase expression or activity promotes AKT activation and enhanced 3D growth of androgen-indifferent mCRPC cells

To begin to investigate the molecular basis of increased tumor growth of Abl kinase-deficient cells, we examined low anchorage growth on a soft 3D collagen matrix, an assay that aligns with the *in vivo* behavior of our mCRPC model [33]. Similar to the tumor growth *in vivo*, ABL/ARG KD cells displayed dramatically enhanced 3D growth compared to vector control cells, with ABL single knockdown producing a larger effect than ARG single knockdown (Fig. 4A). These differences were context specific, and largely not observed under standard tissue culture conditions (Fig. 4B). We found no major changes in focal adhesion kinase (FAK), Src kinase, or ERK kinase activation among the four cell types growing on 3D matrix (Fig. S4). However, Abl-deficient cells displayed a prominent increase in AKT activation, in a pattern that aligned with their 3D growth. ABL/ARG KD cells had the highest level of AKT activation, followed by ABL single knockdown cells and ARG single knockdown cells (Fig. 4C). Quantification of activated pAKT S473 revealed an ∼3.9-fold increase in ABL/ARG KD cells, and ∼2.2-fold and ∼1.4-fold increase in ABL and ARG single knockdown cells respectively, relative to NT/NT vector control cells. Under standard tissue culture conditions, all four cell lines displayed similar levels of AKT activation, consistent with their similar growth under those conditions (Fig. 4D).

**Figure 4:**
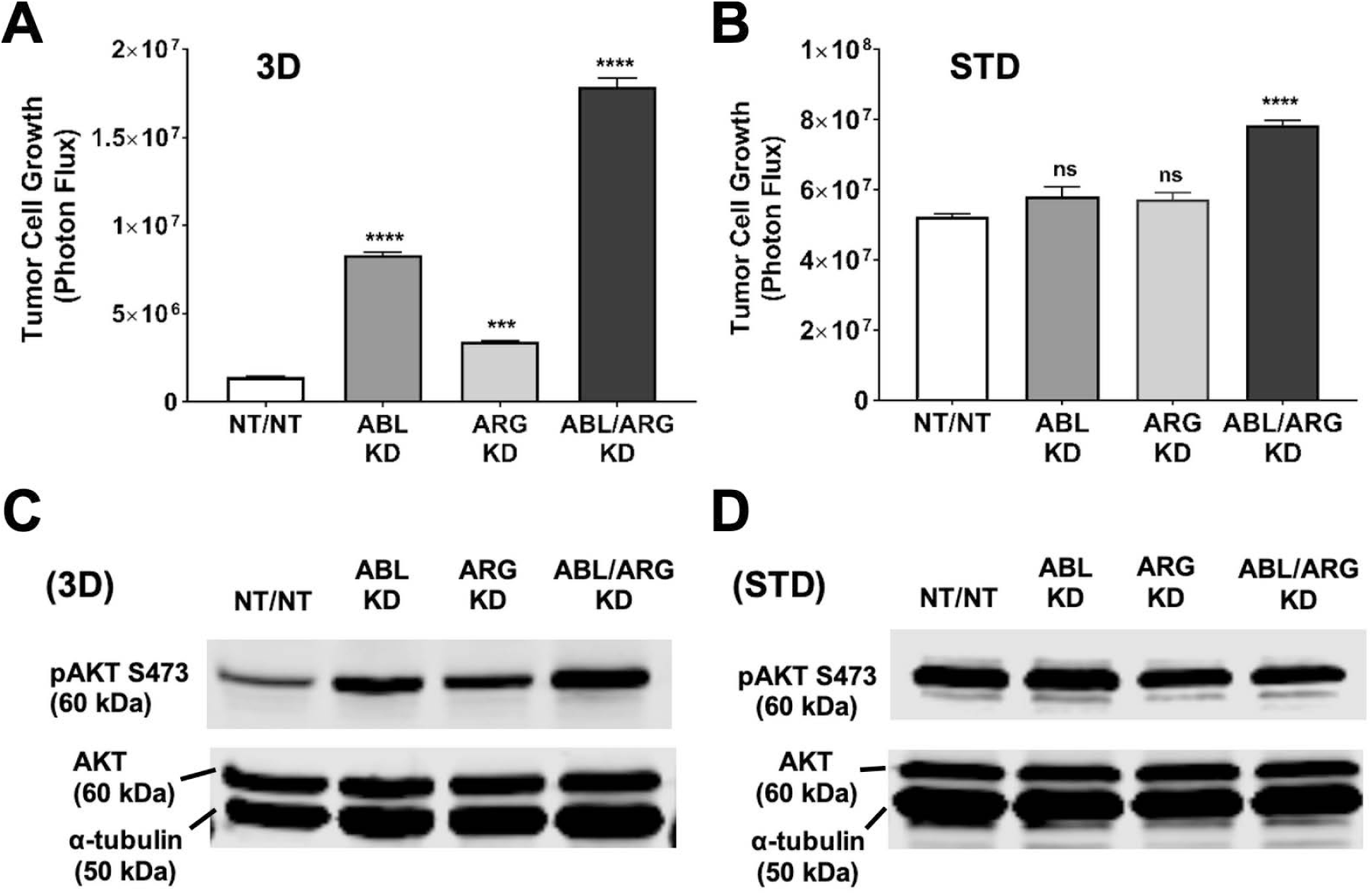
Increased AKT activation and 3D growth of mCRPC cells is associated with Abl kinase deficiency. Graphs of tumor cell growth (photon flux [photons/s]) for Abl family kinase-deficient (ABL KD, ARG KD, ABL/ARG KD) and non-targeting (NT/NT) mCRPC cells growing under **(A)** 3D and **(B)** standard (STD) conditions for seven days. Bars indicate mean +/- SEM for 6 wells/cell type/condition. *** and **** denote statistically significant p-values of < 0.001 and < 0.0001, respectively, and n.s. stands for not significant. One-way ANOVA with Sidak multiple comparison test, α = 0.05. Immunoblot analysis of AKT phosphorylated on serine 473 (pAKT S473), total AKT (AKT), and an α–tubulin loading control for Abl family kinase-deficient and non-targeting mCRPC cells growing under **(C)** 3D and **(D)** STD conditions for 4 days.

We next utilized pharmacological manipulation in conjunction with our 3D growth assay and immunoblot analysis. We predicted that if increased pAKT S473 in ABL KD and ABL/ARG KD cells growing in 3D is a direct result of Abl kinase depletion, then imatinib treatment should produce a large fold change in the growth and phosphorylation of AKT at S473 of non-targeting vector control cells, compared to the fold change for imatinib-treated ABL/ARG KD cells, with much less ABL and ARG protein expression. When we treated Abl family kinase-depleted mCRPC cells growing in 3D with imatinib, we saw significant increases in growth for all cell types relative to their DMSO controls at both 3 uM and 10 uM imatinib, with the higher dose generating an even greater response (Fig. 5A). The fact that ABL/ARG KD cells still displayed increased 3D growth upon treatment with imatinib mostly likely reflects inhibition of residual amounts of ABL (15%) and ARG (20%) remaining in these cells. As anticipated, NT/NT cells experienced the greatest fold change in growth compared to all other cell types at both doses of imatinib, and ABL/ARG KD cells exhibited the smallest fold change in growth, with single knockdown cells displaying an intermediate phenotype (Fig. 5B). In parallel with its ability to promote 3D growth, imatinib treatment dramatically increased phospho-AKT S473 in vector control cells and ARG single knockdown cells, but had minimal to no effect on ABL and ABL/ARG KD cells, in which phospho-AKT levels were already very high (Fig. 5C).

**Figure 5:**
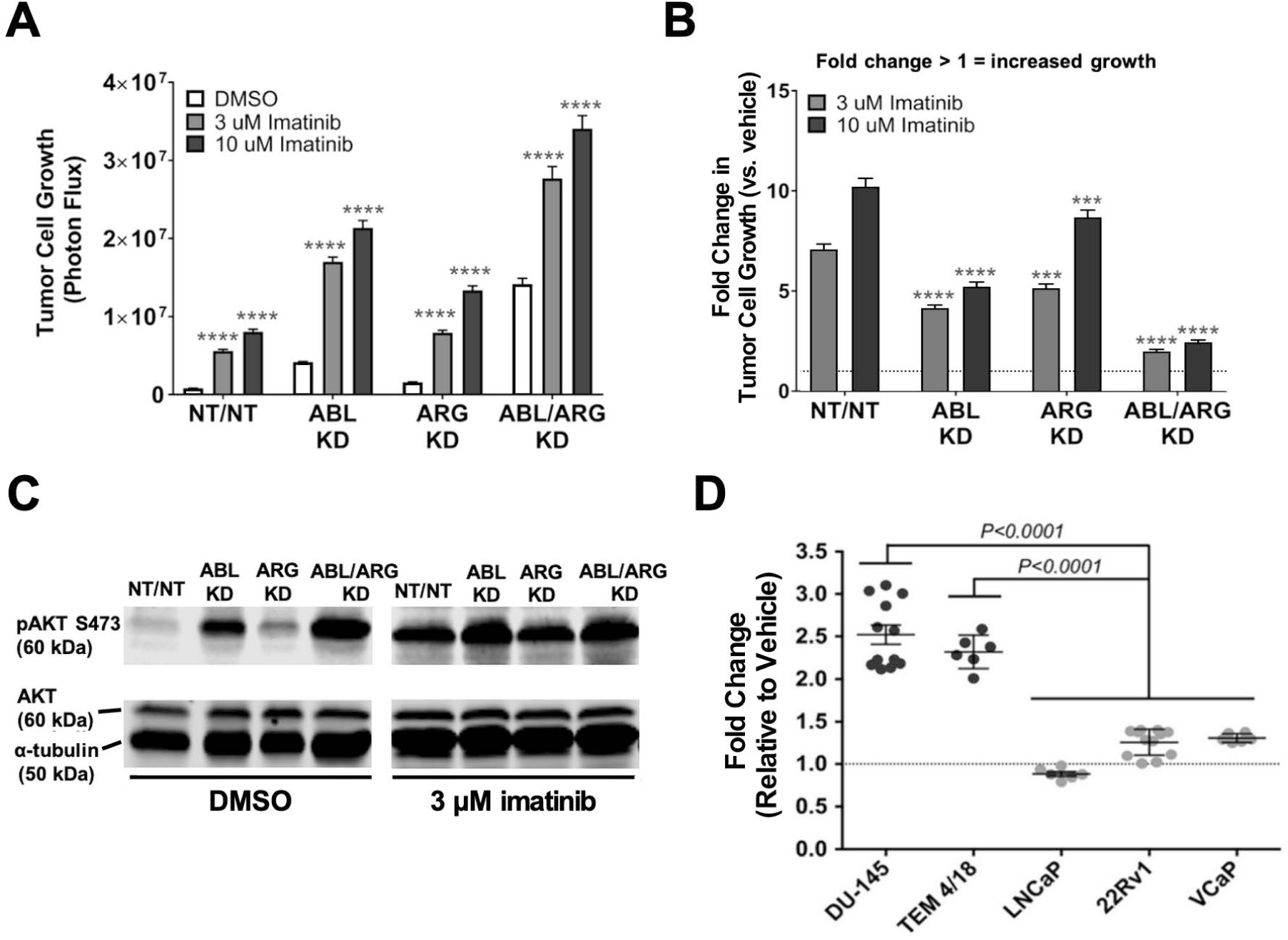
Loss of Abl family kinase activity preferentially enhances the 3D growth of androgen receptor (AR) indifferent prostate tumor cell lines. Graphs of **(A)** tumor cell growth (photon flux [photons/s]) and **(B)** fold change in tumor cell growth relative to vehicle for Abl family kinase-deficient (ABL KD, ARG KD, and ABL/ARG KD) and non-targeting (NT/NT) mCRPC cells growing in 3D and treated with DMSO (white), 3 uM imatinib (light gray), or 10 uM imatinib (black) for 7 days. Bars in A & B indicate mean +/- SEM for 6 wells/cell type/condition. **** in A. indicates statistically significant p-values < 0.0001 within each cell type for a specific dose of imatinib relative to DMSO control. Unpaired t-test with Holm-Sidak correction for multiple t-tests, α = 0.05. *** and **** in B. denote statistically significant p-values of < 0.001 and < 0.0001, respectively, for each Abl family kinase-deficient cell type relative to NT/NT cells for a specific dose of imatinib. Two-way ANOVA with Sidak multiple comparison test, α = 0.05. **(C)** Immunoblot analysis of AKT phosphorylated on serine 473 (pAKT S473), total AKT (AKT), and an α–tubulin loading control for Abl family kinase-deficient and non-targeting mCRPC cells growing in 3D and treated with DMSO or 3 uM imatinib for 4 days. **(D)** Graph of fold change in growth relative to vehicle for AR-indifferent (black) and AR-driven (light gray) prostate cancer cell lines growing in 3D conditions with 3 uM imatinib. Bars indicate mean +/- SEM for 6 wells/cell line. ANOVA with Turkey’s post-test, α = 0.05.

To evaluate the generality of the link between Abl kinase signaling and 3D growth of prostate cancer cells, we tested the effect of imatinib on the 3D growth of 5 additional prostate cancer cell lines. DU145 cells and TEM4-18, another derivative of PC-3 cells [34], are androgen indifferent, while LNCaP, 22Rv1, and VCaP are androgen-responsive cell lines. We found that imatinib promoted the 3D growth of the androgen indifferent cell lines to a significantly greater extent than the androgen responsive cell lines, on which it had little or no effect (Fig. 5D).

To determine the extent to which AKT signaling is required for the increased 3D growth of Abl kinase-deficient cells, we treated cells with increasing doses of the AKT inhibitor MK-2206. MK-2206 abolished the 3D growth of all 4 cell types, even the ABL/ARG KD cells with the highest level of 3D growth (Fig. 6A), and extinguished active AKT in the cells (Fig. 6B). In contrast, under standard culture conditions, the same doses of MK-2206 slowed but did not abolish cell proliferation (Fig. 6C). Together with the previous data, these results indicate that one major consequence of Abl kinase deficiency is to promote AKT signaling, which can result in increased 3D growth of androgen-indifferent prostate cancer cells.

**Figure 6:**
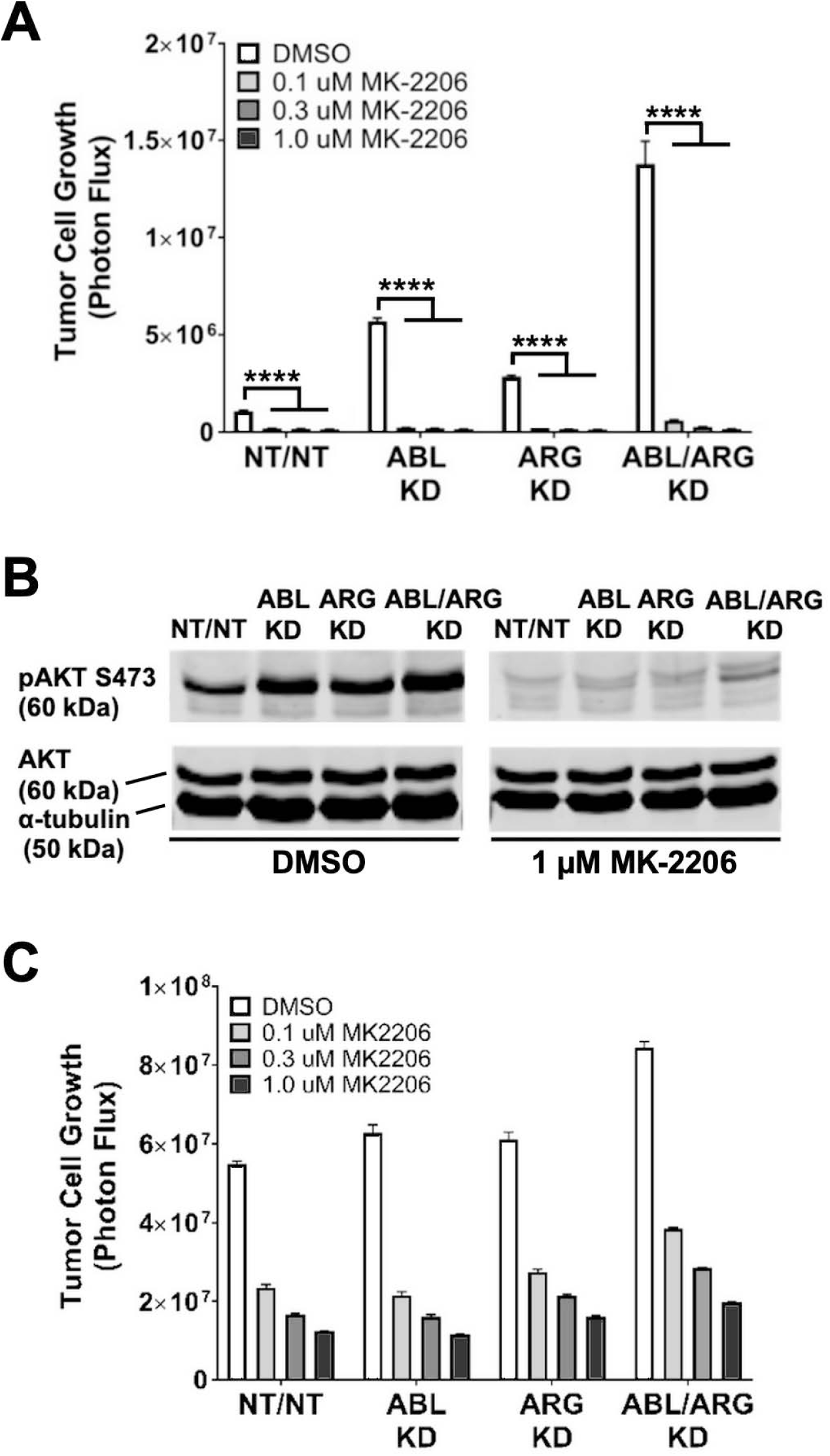
The AKT inhibitor, MK-2206, abolishes the increased 3D growth and AKT activation of Abl family kinase-deficient mCRPC cells. Graphs of tumor cell growth (photon flux in [photons/s]) for Abl family kinase-deficient (ABL KD, ARG KD, ABL/ARG KD) and non-targeting (NT/NT) mCRPC cells under 3D **(A)** and standard **(C)** conditions and treated with DMSO (white), 0.1 uM MK-2206 (light gray), 0.3 uM MK-2206 (dark gray), or 1.0 uM MK-2206 (black) for 7 days. Bars in A & C indicate mean +/- SEM for 6 wells/cell type/condition. **** in A signifies statistically significant p-values <0.0001. Unpaired t-test with Holm-Sidak correction for multiple t-tests, α = 0.05. **(B)** Immunoblot analysis of AKT phosphorylated on serine 473 (pAKT S473), total AKT (AKT), and an α–tubulin loading control for Abl family kinase-deficient and non-targeting mCRPC cells growing in 3D for 4 days and treated with DMSO or 1.0 uM MK-2206 for 1 day.

### Reverse phase protein array (RPPA) analysis reveals upstream and downstream effectors of AKT signaling in Abl kinase-deficient mCRPC cells

To investigate the link between Abl family kinases and AKT signaling we generated lysates of NT/NT and ABL/ARG KD cells growing in 3D conditions for analysis by RPPA using 304 antibodies to cancer-relevant signaling proteins. An arbitrary cut off was set for an average fold change of 1.2 or 0.8. Antibodies with average fold changes meeting the cutoff criteria were visualized by a heatmap using median centered normalized log_2_ values (Fig. 7A; see Table S1 for full results of the RPPA).

**Figure 7:**
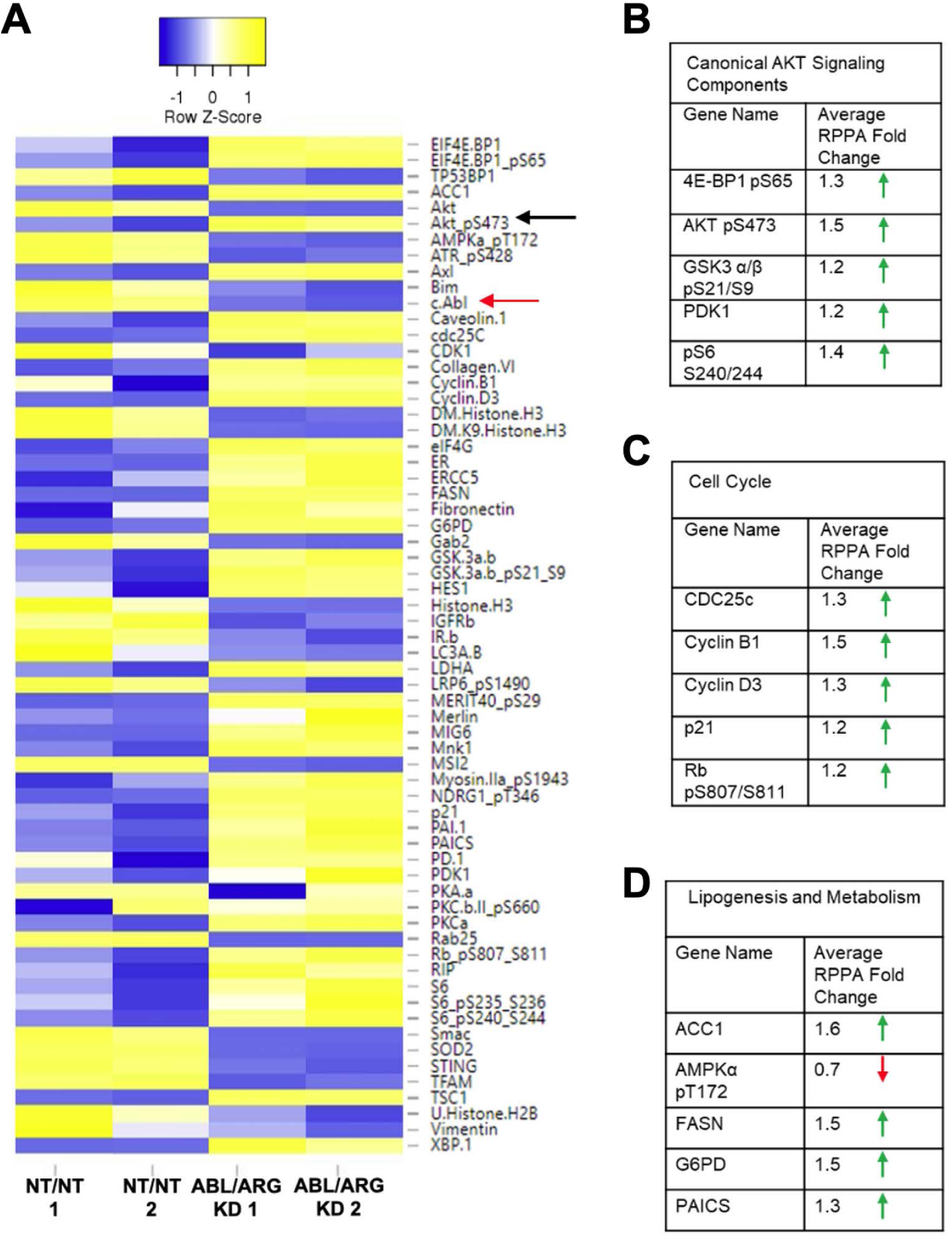
Reverse phase protein array (RPPA) analysis of Abl family kinase-deficient mCRPC cells growing in 3D. **(A**) Heatmap displaying median-centered, normalized log_2_ values of differentially expressed antigens (DEAs). Lysates of NT/NT and ABL/ARG KD mCRPC cells grown under 3D conditions for 4 days were analyzed by RPPA. Antibodies with average normalized linear fold changes > 1.2 or < 0.8 were considered DEAs. Rows corresponding to pAKT S473 (black) and c-Abl (red) are marked with arrows. Tables of average normalized linear fold changes for selected groupings of DEAs: **(B)** AKT signaling components, **(C)** cell cycle factors, and **(D)** genes involved in lipogenesis and metabolism.

As expected, pAKT S473 (fold change = 1.5; black arrow Fig. 7A) and c-ABL (fold change = 0.7; red arrow Fig. 7A) were among the list of 64 differentially expressed antigens (DEAs). Additionally, several canonical AKT signaling components were differentially expressed (selected list in Fig. 7B). Pathway analysis with DAVID showed that 26.3% of DEAs were associated with the PI3K-AKT signaling pathway (P = 6.7×10^−8^). Cell cycle factors were also greatly enriched in DEAs (selected list in Fig. 7C), with DAVID pathway analysis showing 15.8% being associated with the cell cycle (P = 2.2×10^−6^). Proteins involved in metabolism and lipogenesis (selected list in Fig. 7D) were also prevalent among DEAs and 15.8% of them were associated with AMPK signaling (DAVID pathway analysis; P = 2.2×10^−6^).

We further validated our RPPA data by immunoblotting proteins from each of the categories identified during DAVID pathway analysis (Fig. S5). Expression of the canonical AKT signaling target S6 Ribosomal Protein phosphorylated at serine 240/244 (pS6 S240/244) (Fig. S5A, D), and cyclin D3 (Fig. S5B, E), mirrored the pattern of pAKT S473 expression in Abl family kinase-depleted cells, with the highest expression levels of these antigens in ABL/ARG KD double knockdown and ABL KD single knockdown cells. Conversely, expression of AMPKα phosphorylated at its activating threonine172, a site negatively regulated by AKT [46], was downregulated in ABL/ARG KD cells (Fig. S5C, F).

AXL, a receptor tyrosine kinase and known activator of PI3K/AKT signaling [47], was also among the DEAs identified in our RPPA dataset. Therefore, we assessed expression of total AXL, as well as expression of an autophosphorylation site at tyrosine 779 (pAXL Y779) known to bind the p85α and p85β subunits of PI3K [47]. Both phospho-AXL and total AXL levels were increased in the ABL single knockdown and ABL/ARG double knockdown cells relative to NT/NT vector control cells. Quantification revealed an ∼1.2 fold increase pAXL Y779 in ABL single knockdown cells and ∼1.4 fold increase in ABL/ARG double knockdown cells, while expression of total AXL was substantially higher in ABL KD cells (∼2.3 fold increase) and ABL/ARG KD cells (∼3.3 fold increase) (Fig. 8A). Moreover, pharmacological inhibition of Abl kinase activity in NT/NT cells using imatinib led to an ∼1.4-fold increase in total AXL expression in cells growing on 3D matrix (Fig. S6).

**Figure 8:**
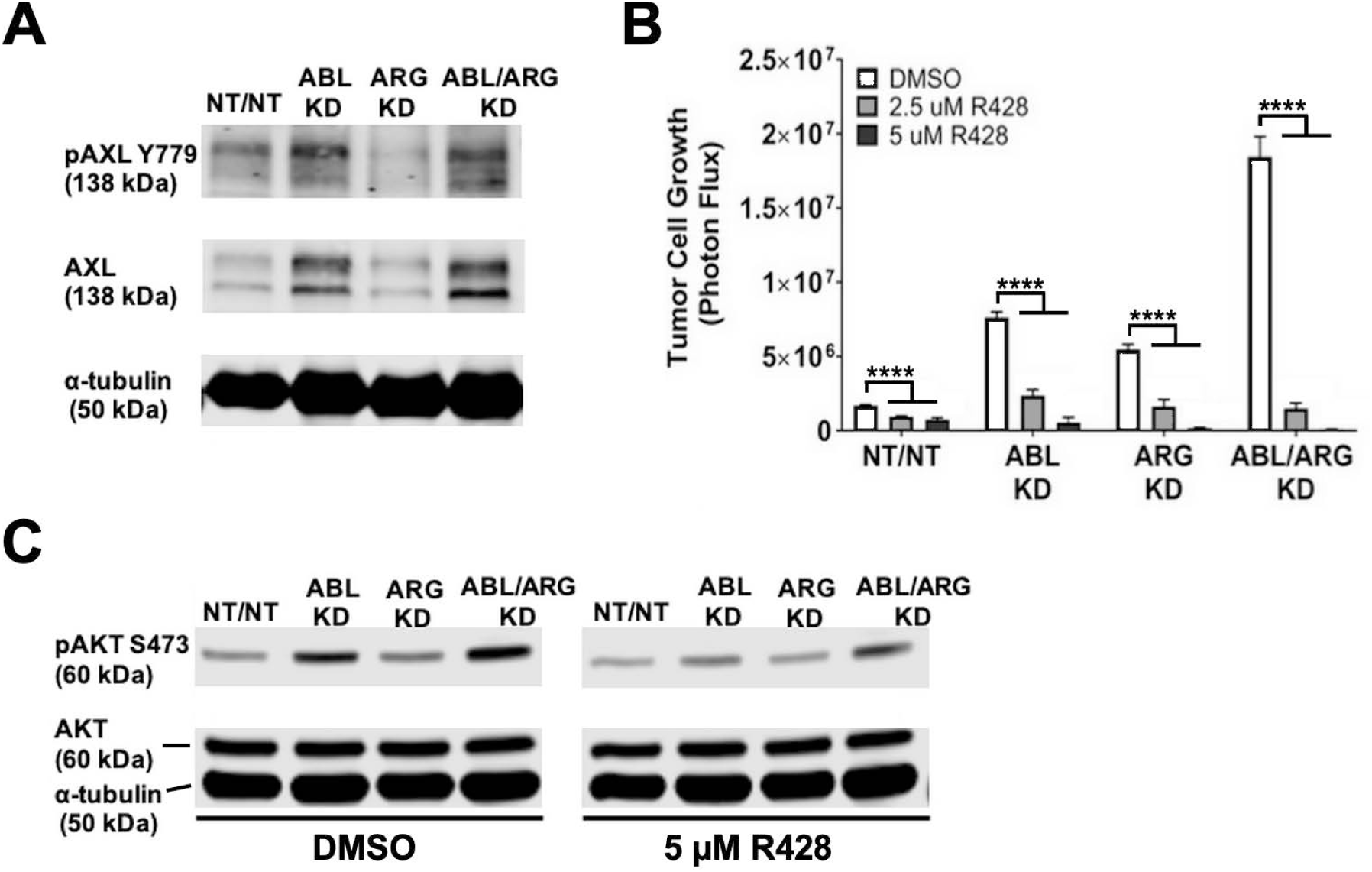
AXL activation and expression is associated with the 3D growth phenotype and AKT hyperactivity of Abl family kinase-deficient mCRPC cells. **(A)** Immunoblot analysis of AXL autophosphorylated on tyrosine 779 (pAXL Y779), total AXL (AXL), and α–tubulin loading control for Abl family kinase-deficient (ABL KD, ARG KD, ABL/ARG KD) and non-targeting (NT/NT) mCRPC cells growing in 3D for 4 days. **(B)** Graph of tumor cell growth (photon flux [photons/s]) of Abl family kinase-deficient and non-targeting mCRPC cells growing in 3D and treated with DMSO (white), 2.5 uM (gray), or 5 uM (black) of the specific AXL inhibitor, R428 for 7 days. Bars indicate mean +/- SEM for 6 wells/cell type/condition. **** indicates statistically significant p-values ≤ 0.0001. Unpaired t-test with Holm-Sidak correction for multiple t-tests, α = 0.05. **(C)** Immunoblot analysis of AKT phosphorylated on serine 473 (pAKT S473), total AKT (AKT), and an α–tubulin loading control for Abl family kinase-deficient and non-targeting mCRPC cells growing in 3D for 4 days and treated with DMSO or 5 uM R428 for 1 day.

To investigate whether increased AXL expression in Abl-deficient cells contributes to AKT activation and 3D growth, we treated cells with the specific AXL inhibitor, R428 (BGB324) [48]. R428 strongly suppressed the 3D growth of Abl-deficient cells (Fig. 8B) and partially canceled the increased AKT activation seen in ABL kinase single knockdown and ABL/ARG double knockdown cells (Fig. 8C). Taken together, results of our RPPA analysis and subsequent validation studies solidify AKT activation as a major pro-tumor outcome of Abl kinase deficiency and identify increased AXL expression as a candidate mediator of increased AKT signaling and 3D growth of Abl kinase-deficient cells.

## Discussion

A major finding of our study was the dramatic increase in the rate of tumor progression and metastasis of Abl kinase-deficient tumor cells, establishing that Abl kinases can function as suppressors of progression and metastasis in a pre-clinical model of mCRPC. Interestingly, in the TRAMP mouse model of prostate cancer, treatment with the Abl kinase inhibitor imatinib resulted in the selective outgrowth of neuroendocrine-like tumors [49], and imatinib failed in mCRPC clinical trials, where it was associated with a lower probability of PSA decline and reduced overall and progression-free survival [50]. Both of these observations are consistent with progression and metastasis suppressor functions for Abl kinases in mCRPC.

Although inhibiting Abl kinase activity has been advanced as potential therapeutic strategy for treating solid tumors [22], our new results add to a growing number of studies that point to the ability of Abl kinases to suppress cell motility, promote apoptosis, and negatively regulate cell proliferation and tumor progression in multiple settings [23, 26-29, 32, 51-55]. Anti-tumor activity of Abl kinases has been linked to signaling by tumor suppressor EphB4 [29], p53-P21 signaling [23, 27, 55], negative regulation of Ras-MAP kinase signaling [26], and conversion of the proto-oncogenic YAP1 transcription factor from a pro-growth to a pro-apoptotic function upon DNA damage [54].

Our new data point to additional anti-tumor functions for Abl kinases that are distinct from those that have previously been described. We find that Abl family kinase depletion results in a dramatic upregulation of AKT pathway signaling in our p53-null, PTEN-deficient mCRPC model, with no obvious impact on signaling via the MAP kinase pathway (Figure S4). Pre-clinical data link hyperactivation of the AKT pathway to the emergence of mCRPC [16-20], but our new data suggest that AKT signaling can continue to sustain the 3D growth of fully AR-indifferent mCRPC cells. Moreover, our data support the view that even in the context of PTEN deficiency other factors, such as Abl family kinase signaling, can continue to restrain the full oncogenic capacity of the AKT pathway.

The mechanism by which Abl kinase deficiency results in upregulation of the AKT pathway requires further study. However, our RPPA data nominated increased protein expression of the receptor tyrosine kinase AXL as one major input into the increased AKT signaling in Abl-deficient cells. In prostate cancer, AXL signaling has been associated with metformin, taxotere, and docetaxel resistance [56-58], as well as EMT-like events, increased migration and invasion, proliferation and tumor growth [59-61]. Conversely, some studies have reported anti-proliferative effects for AXL, which may be related to its ability to induce tumor cell dormancy in some settings [58, 62]. AXL has also recently emerged as a direct target of tumor suppressive micro-RNAs in prostate cancer models [63, 64].

The new findings presented here have important implications for ongoing clinical efforts to target AKT in mCRPC, such as the phase III trial combining AKT inhibitor ipatasertib with abiraterone (NCT03072238). The scientific rationale for this combination trial involves reciprocal feedback between androgen receptor signaling and PI 3-kinase/Akt signaling in PTEN-deficient prostate cancer [21], such that inhibiting one pathway activates the other, suggesting dual inhibition as a strategy. However, this clinical trial is not designed to investigate efficacy related to the presence or absence of AR-indifferent and neuroendocrine-like disease and also will not address the potential utility of AKT inhibition in mCRPC that has already progressed to an AR-indifferent state. Our data suggest the possibility that tumors that evolve resistance to AR inhibitors such as abiraterone yet remain AR-driven may be less amenable to AKT inhibitors than tumors that evolve to an AR-indifferent state. Important future directions based on results presented here include testing the efficacy of AKT and AXL inhibitors in pre-clinical *in vivo* models of AR-indifferent and AR-driven metastatic prostate cancer and investigating at the protein level AXL and AKT expression and markers of Abl kinase activity in mCRPC clinical specimens. It will also be important to investigate the mechanism by which Abl kinase deficiency results in upregulation of AXL protein and the extent to which factors in addition to AXL expression contribute to AKT pathway hyperactivation in Abl deficient cells. Such investigations 8 may yield additional treatment strategies for AR-indifferent, NE-like prostate cancers, especially 9 those with PTEN deficiency, which represents ∼50-60% of all NE-like mCRPC [13].

## Acknowledgements

Dr. Brown is the current University of Iowa Andersen-Hebbeln Professor in Prostate Cancer Research. This endowed professorship provided financial support related to this research effort. Core facilities used in these studies were supported by NIH grant P30 CA086862.

