## Supplemental Figures for "Abl kinase deficiency promotes AKT pathway activation and prostate cancer progression and metastasis"

### Supplemental Figure Legends

**Fig. S1 Expression of neuroendocrine markers in PC-3 vs. GS689.Li cells**

(**A & B**) chromogranin A (CHGA) and DAPI staining. (**C&D**) Rb staining. (**E&F**) E-cadherin and FOXA2 staining. Arrows indicate E-cadherin negative, FOXA2-positive cells in the parental PC-3 population. GS689.Li cells are uniformly E-cadherin-negative and FOXA2-positive. Bars = 15 um.

**Figure S2: Post-implantation analysis of ABL and ARG protein expression**

Immunoblot analysis of ABL and ARG proteins in mCRPC cells recovered from primary tumors of mice bearing Abl family kinase-deficient (ABL KD, ARG KD, and ABL/ARG KD) and non-targeting (NT/NT) tumors for 3 mice/tumor type. Values beneath ABL and ARG protein bands represent blot intensity values normalized by an α–tubulin loading control. Values beneath α–tubulin protein bands represent raw blot intensity values.

**Figure S3: Abl family kinase-deficient mCRPC cells display distinct morphological phenotypes during migration**

The morphology of Abl family kinase deficient (ABL KD, ARG KD, and ABL/ARG KD) and non-targeting (NT/NT) mCRPC cells migrating on 2D collagen I was monitored via time-lapse microscopy and analyzed at t = 20 minutes. Time-lapse micrographs of **(A)** NT/NT, **(B)** ABL KD, **(C)** ARG KD, and **(D)** ABL/ARG KD mCRPC cell populations. Insets feature lamellae (L) and morphology of representative cells. Scale bar = 50 μm. Graphs depicting **(E)** the average aspect ratio and **(F)** average roundedness for Abl family kinase-deficient and non-targeting mCRPC cells. * and ** in E and F denote statistically significant p-values of $< 0.05$ and $< 0.01$, respectively. One-way ANOVA w/ Dunnett's multiple comparison, n = 27 cells, $\alpha = 0.05$.

**Figure S4: The increased 3D growth of Abl family kinase-deficient mCRPC cells is not associated with activation of FAK, SRC, or ERK**

Immunoblot analysis of FAK phosphorylated at tyrosine 397 (pFAK Y397), total FAK, SRC phosphorylated at tyrosine 416 (pSRC Y416), total SRC, ERK 1/2 phosphorylated at threonine 202 and tyrosine 204 (pERK T202/Y204), and total ERK 1/2 for Abl family kinase deficient (ABL KD, ARG KD, and ABL/ARG KD) and non-targeting (NT/NT) mCRPC cells growing under 3D conditions for 4 days.

**Figure S5: Western blot validation of select RPPA differentially expressed antigens (DEAs)**

Immunoblot analysis of **(A)** S6 Ribosomal Protein phosphorylated on serine 240/244 (pS6 S240/244), **(B)** Cyclin D3, **(C)** AMPKα phosphorylated on threonine 172 (pAMKα T172), and their respective α–tubulin loading controls for Abl family kinase-deficient (ABL KD, ARG KD, and ABL/ARG KD) and non-targeting (NT/NT) mCRPC cells growing in 3D for 4 days. Values beneath pS6 S240/244, Cyclin D3, and pAMKα T172 protein bands represent blot intensity values normalized by an α–tubulin loading control. Values beneath α–tubulin protein bands represent raw blot intensity values. Tables summarizing normalized fold changes for ABL KD, ARG KD, and ABL/ARG KD mCRPC cells relative to NT/NT for **(D)** pS6 S240/244, **(E)** Cyclin D3, and **(F)** AMPKα T172.

**Figure S6: Loss of Abl family kinase activity results in increased AXL expression**

**(A)** Immunoblot analysis of total AXL in non-targeting (NT/NT) mCRPC cells growing in 3D and treated with DMSO or 3 uM imatinib for 4 days. Values beneath total AXL protein bands represent blot intensity values normalized by an α–tubulin loading control. Values beneath α–tubulin protein bands represent raw blot intensity values. **(B)** Table summarizing % increase or decrease in total AXL expression calculated from normalized values in A.

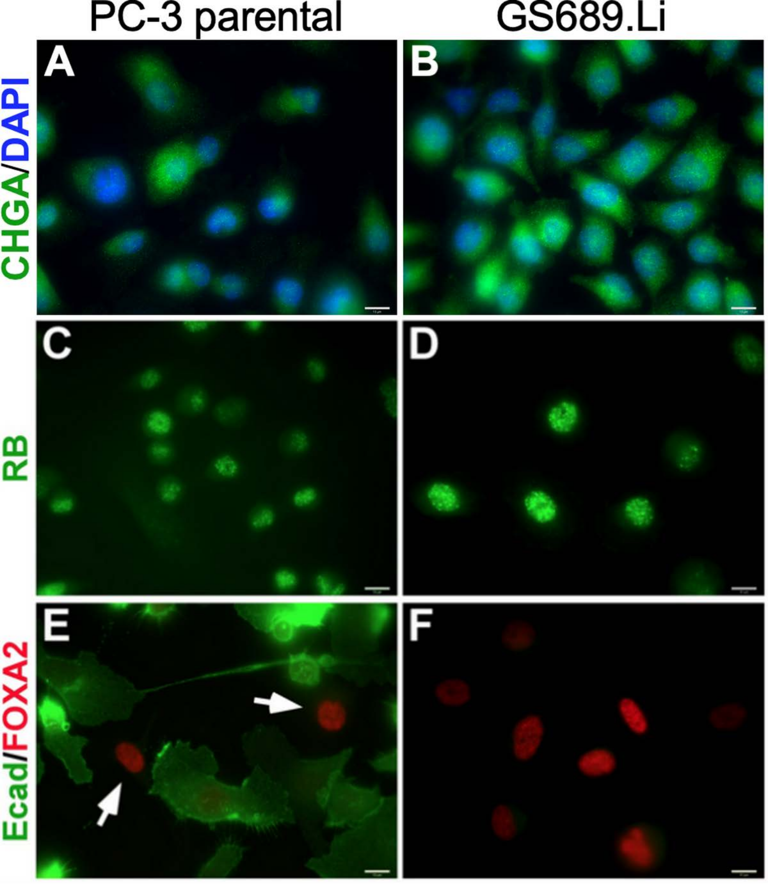

PC-3 parental
GS689.Li
CHGA/DAPI
RB
Ecad/FOXA2
A
B
C
D
E
F

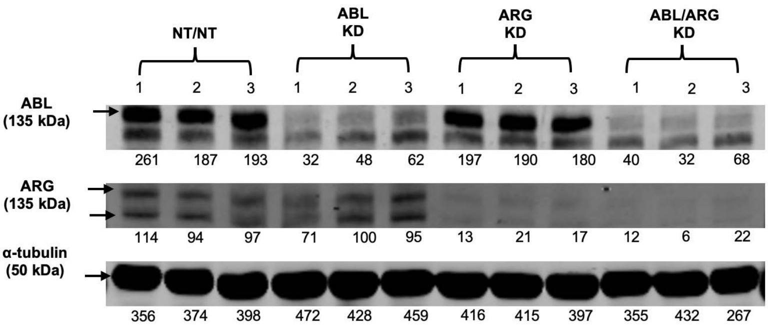
NT/NT
ABL KD
ARG KD
ABL/ARG KD
1 2 3 1 2 3 1 2 3 1 2 3
ABL (135 kDa)
261 187 193 32 48 62 197 190 180 40 32 68
ARG (135 kDa)
114 94 97 71 100 95 13 21 17 12 6 22
α-tubulin (50 kDa)
356 374 398 472 428 459 416 415 397 355 432 267

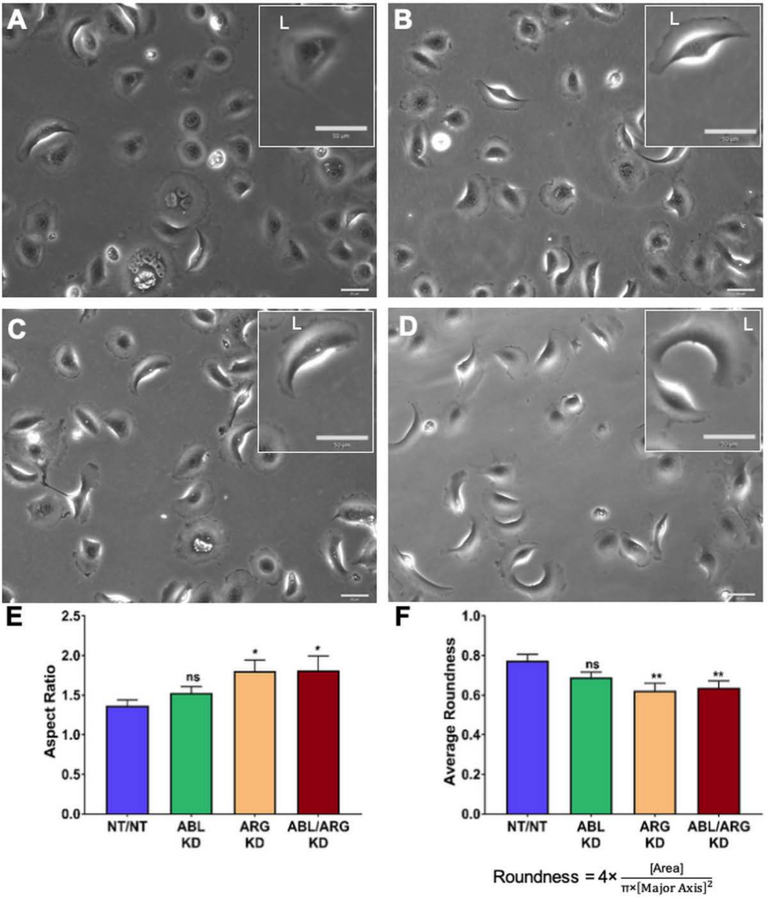
A
L
B
L
C
L
D
L
E
Aspect Ratio
2.5
2.0
1.5
1.0
0.5
0.0
ns
*
*
NT/NT
ABL KD
ARG KD
ABL/ARG KD
F
Average Roundness
1.0
0.8
0.6
0.4
0.2
0.0
ns
**
**
NT/NT
ABL KD
ARG KD
ABL/ARG KD
Roundness = 4× [Area] / π×[Major Axis]²

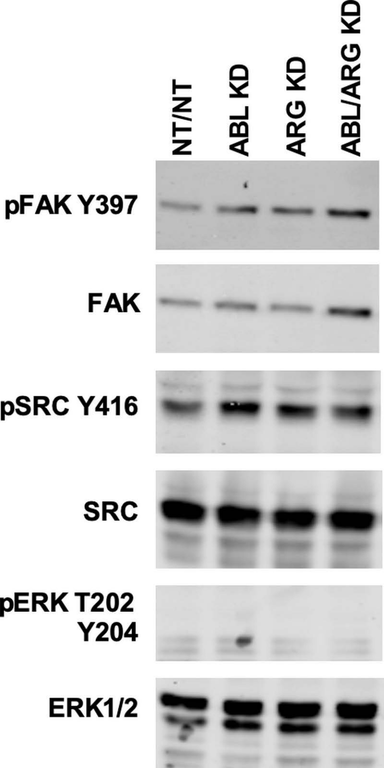
NT/NT
ABL KD
ARG KD
ABL/ARG KD
pFAK Y397
FAK
pSRC Y416
SRC
pERK T202
Y204
ERK1/2

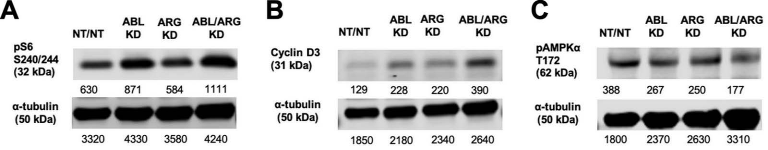


**D**

| pS6 S240/244 | |
|---|---|
| Genotype | Normalized fold change relative to NT/NT |
| ABL KD | 1.7 ↑ |
| ARG KD | 1.2 ↑ |
| ABL/ARG KD | 1.7 ↑ |

**E**

| Cyclin D3 | |
|---|---|
| Genotype | Normalized fold change relative to NT/NT |
| ABL KD | 1.8 ↑ |
| ARG KD | 1.7 ↑ |
| ABL/ARG KD | 3.0 ↑ |

**F**

| pAMPKα T172 | |
|---|---|
| Genotype | Normalized fold change relative to NT/NT |
| ABL KD | 0.7 ↓ |
| ARG KD | 0.6 ↓ |
| ABL/ARG KD | 0.5 ↓ |

A

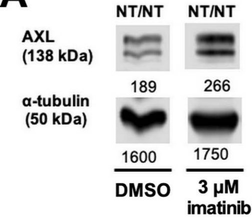


B

| Genotype | % increase or decrease AXL 3 μM imatinib vs. DMSO |
|---|---|
| NT/NT | 41% ↑ |
